# The Coronavirus Network Explorer: Mining a large-scale knowledge graph for effects of SARS-CoV-2 on host cell function

**DOI:** 10.1101/2020.09.14.296327

**Authors:** Andreas Krämer, Jean-Noël Billaud, Stuart Tugendreich, Dan Shiftman, Martin Jones, Jeff Green

**Affiliations:** QIAGEN Digital Insights, Redwood City, CA 94063, USA

## Abstract

Building on recent work that identified human host proteins that interact with SARS-CoV-2 viral proteins in the context of an affinity-purification mass spectrometry screen, we use a machine learning-based approach to connect the viral proteins to relevant biological functions and diseases in a large-scale knowledge graph derived from the biomedical literature. Our aim is to explore how SARS-CoV-2 could interfere with various host cell functions, and also to identify additional drug targets amongst the host genes that could potentially be modulated against COVID-19. Results are presented in the form of interactive network visualizations, that allow exploration of underlying experimental evidence. A selection of networks is discussed in the context of recent clinical observations.

## Introduction

Severe acute respiratory syndrome coronavirus 2 (SARS-CoV-2), a member of the coronavirus family, is the etiologic agent of the pandemic COVID-19. Like other positive-stranded RNA viruses, its encoded proteins interact with proteins of the infected host cell at various stages of the replicative cycle, including with those involved in the immune response. Such proteins therefore represent possible targets for the development of antiviral strategies. Recently Gordon *et al*. (1) identified human host proteins that bind to overexpressed SARS-CoV-2 viral proteins in immortalized human cells using an affinity-purification mass spectrometry screen. Proteins identified were functionally characterized and screened for existing drug targets to identify drugs that could potentially be repurposed against COVID-19 (1).

Our goal is to illuminate possible molecular mechanisms permitting viral proteins to affect a range of host cell and immune functions that have been shown to be relevant in the context of COVID-19. For this purpose we integrate SARS-CoV-2 viral proteins into a large-scale knowledge graph (KG) through the viral-host protein interactions identified in Gordon *et al*. (1). The KG represents existing experimental evidence from the biomedical literature mostly in the form of signed cause-effect relationships (edges). These edges connect different entities (nodes) including genes, drugs, microRNAs, biological functions, diseases, and pathways; and they relate to different underlying functional mechanisms such as expression/transcription, activation/inhibition, phosphorylation, protein-protein binding among others. Each relationship has a positive or negative sign indicating the direction of the effect, i.e. whether it leads to an increase or decrease (2). Underlying the KG is the QIAGEN Knowledge Base (KB), a large structured collection of biomedical content that has been manually curated from the literature for the past 20 years, and also integrates content from third-party databases. At present, the KB contains over 7 million findings, a large subset of which are represented in the KG with approximately 120,000 nodes and 3.7 million edges.

## Approach

The outline of our approach is shown in Figure 1. For each member of a pre-selected set of biologically relevant endpoint functions (including some diseases and signaling pathways), we construct a network (a subgraph of the KG) that connects viral proteins to the specified endpoint through causal paths. These causal paths involve genes (or other molecules) that are important in the given functional context, and therefore indicate a possible mechanism with which viral proteins may affect the particular host function. Because of the high connectivity of the KG, a large number of molecules may be associated with the host function, so in order to construct focused networks, care must be taken regarding the choice of intermediary genes and molecules. For this reason, we use a machine learning model trained on prior knowledge, that computes a signed score for each gene indicating the likelihood of an activating or inhibiting effect on the endpoint function, where the direction of the effect is given by the sign of the score. The purpose of the model is twofold: (1) the prediction of novel gene-function associations independent of whether direct literature evidence is available or not, and (2) the prioritization of genes predicted to be most relevant in the given functional context.

**Fig. 1.**
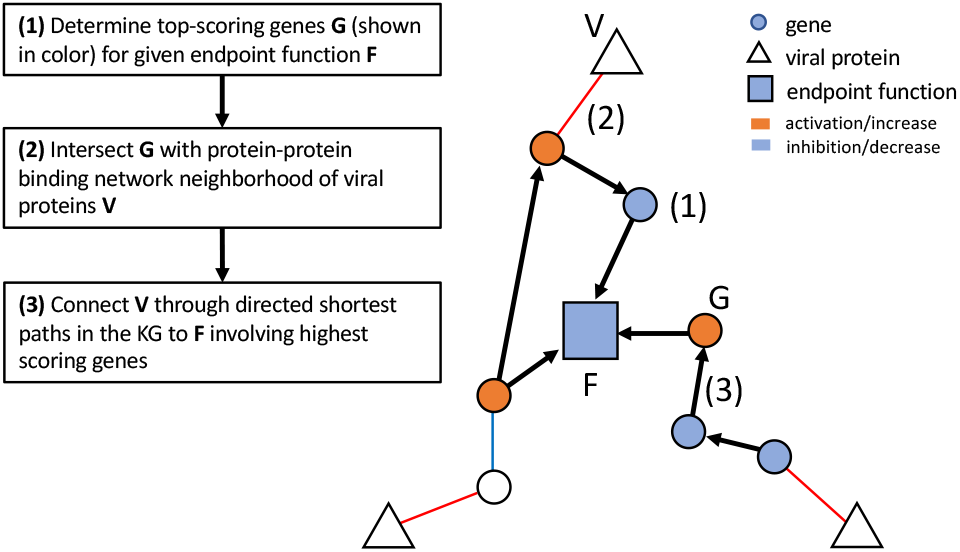
High-level outline of the network algorithm (see also Methods section). Network creation involves three major steps: (1) the selection of top-scoring genes in the context of the given endpoint function as described in the Methods section and shown in Figure 2, (2) the intersection of this set of genes with network neighborhoods of viral proteins, and (3) the connection of viral proteins to the endpoint function through shortest paths in the knowledge graph.

Connections from the viral proteins to the selected endpoint functions were made through short directed paths in the KG involving the highest scoring genes in the given context, where protein-protein binding edges can be traversed in both ways. Therefore, in every network there will always be at least one causal path from any viral protein to the endpoint function through edges supported by literature evidence. Since viral proteins are connected through proteinprotein binding edges to the rest of the network it is impossible to make predictions about their direction of effect on the endpoint function. However, we can use other available information to make inferences about the activation status of the intermediary genes. For instance, recent research (3–7) indicates that SARS-Cov-2 has an activating effect on coagulation of blood (severe coagulopathy has been detected in many patients with severe disease) which allows us to infer that network proteins predicted to have an activating effect on coagulation would potentially need to be inhibited to counteract viral effects, i.e. drugs targeting those proteins would need to be antagonists. Many of the intermediary genes found in our networks are already targets for existing drugs that could potentially be repurposed against COVID-19. Considering the direction of the effect is an important criterion for the selection of most promising candidates.

Existing network-based approaches in the context of COVID-19 mostly focus on protein-protein and co-expression networks using standard enrichment-based functional and pathway analysis, or employing network topology measures to identify possible targets (8–12). Gysi *et al*. (13), based on the Gordon *et al*. (1) dataset in conjunction with the human interactome, provide a systematic exploration of state-of-the art network algorithms (proximity-based, diffusion-based, and based on graph convolutional networks) with the primary goal of ranking candidate drugs for repurposing. To our knowledge, none of these approaches currently leverages the direction of effects, i.e. activation vs. inhibition.

## Implementation

We implemented a web-based interface through which networks computed with the algorithm outlined above can be accessed for interactive exploration. This interface enables the user to select a particular endpoint function, as well as the direction of the effect (activating or inhibiting) that infection with SARS-CoV-2 is thought to have on this function. In the resulting displayed networks (for examples see Figures 3-5) the user can click on any entity (node or edge) to reveal underlying findings from the literature, or a description of the selected entity itself. Colors of gene nodes indicate whether that particular gene is inferred to be activated (orange) or inhibited (blue) in order to achieve the preselected effect on the endpoint function. In addition, we also annotated genes with associated signaling pathways, colored according to their predicted activation state. Nodes whose activation status cannot be inferred, either because they are only connected by unsigned binding edges, or do not meet a required score threshold, are shown in white. Genes that are targets for existing drugs are marked with a purple border. Selecting such a node will display the drugs targeting that gene along with each drug’s predicted effect on the endpoint function. The web application also allows the user to search for specific genes or pathways in order to narrow down the number of networks to peruse.

## Methods

### Prior knowledge-based machine learning model for scoring gene-function relationships

Prediction and scoring of causal gene-function relationships involves two parts, an unsupervised step to construct gene embeddings in a high-dimensional vector space, and a supervised step to make predictions about the gene-function relationships themselves. Building on an assumption that expression relationships encode information about gene function, the unsupervised step builds gene feature vectors by leveraging downstream causal gene expression signatures derived from the literature. It is important to distinguish this from gene expression patterns found in expression datasets: Here, expression signatures are created from individual published gene-to-gene expression or transcription relationships, not from datasets. The supervised step then employs a linear least-square regression model for each function using observed signed causal gene-function relationships from the literature as training data. Below we provide a brief description of the steps involved in this method, a deeper discussion and evaluation of the approach will be given elsewhere (manuscript in preparation).

Causal gene expression signatures are represented by a signed bipartite adjacency matrix *W* where rows represent the genes (or other molecules) for which we wish to compute embeddings, and columns represent genes of the downstream expression signature (Figure 2a). The matrix elements are 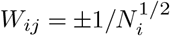 if an observation of gene *i* regulating gene *j* exists in the literature, where the positive sign stands for observed up-regulation and the negative sign for observed down-regulation. It is *W_i_j* = 0 if there is no recorded relationship, and *N_i_* denotes the total number of genes found to be regulated by gene *i*. From *W* we construct the “co-activity” matrix

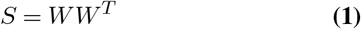

which is positive definite with diagonal elements being equal to one. We then take the low-rank approximation of *S*,

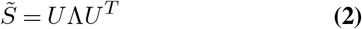

where Λ is an *N*-dimensional diagonal matrix containing the top *N* eigenvalues of *S*, and column vectors of *U* are the corresponding *N* normalized eigenvectors. The row vectors of *U* form the desired *N*-dimensional feature representations of genes (Figure 2b). In the model used for this application we included 2314 genes, and set *N* = 100.

Causal gene-function effects from the literature are captured similarly in a signed bipartite adjacency matrix *Y*, with rows representing genes, and columns representing downstream functions (Figure 2c). It is *Y_ij_* = ±1 if the gene-function effect is activating (+1), or inhibiting (−1), else *Y_ij_* =0 if there is no observed effect. We build a model predicting the effect of gene *i* on function *j* from the gene feature vector *x_i_* by using linear regression, i.e. minimizing the least-squares loss

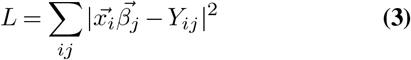

where the parameter vector 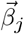 is determined for each function *j*. It follows that

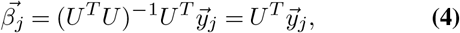

so that predicted signed scores *P_ij_* can then be expressed as orthogonal projections of the column vectors 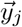 of *Y* onto the subspace spanned by the top eigenvectors of *S* (Figure 2d),

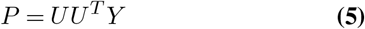

**Fig. 2.**
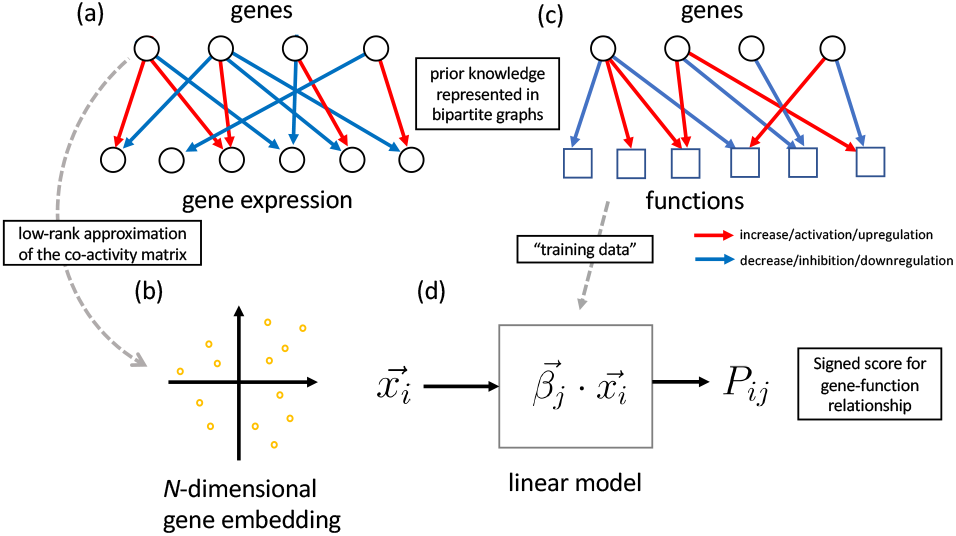
Outline of the prior knowledge-based machine learning approach for scoring gene-function relationships as described in the Methods section. The algorithm assumes that gene expression changes caused by a given gene encode its effect on biological function. *N*-dimensional embeddings are computed for each gene from literature-derived gene expression relationships. These in turn are used as feature vectors in a linear regression model trained on gene-function effects from the literature.

In order to determine top-scoring genes for a given function *j*, we sort the elements of row vectors 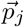 of *P* by their absolute value. A meaningful cutoff is obtained by looking at the distribution of values in 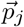, i.e. applying a transformatin to *z*-scores,

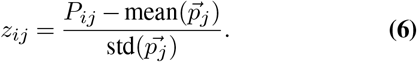

### Network algorithm

The network-generating algorithm consists of the following steps (see Figure 1):

For each endpoint function *F* we determine the set of top-scoring genes associated with *F* where each gene *g* meets acut off |*z_gF_*| ≥ 2. Then, for each viral protein *V* we construct the network neighborhood of *V* by collecting all genes directly connected to *V* (through edges from Gordon *et al*. (1)), and additionally genes that can be reached from *V* by traversing two subsequent protein-protein binding edges (“2 hops”) but also obey a “specificity” criterion ensuring that connections do not occur through network hubs, and providing an external parameter to control network size. In particular, for a given gene *g* that is connected through 2 hops to *V* we calculate the quantity

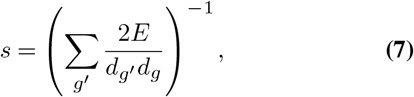

where *E* is the total number of protein-protein binding edges, *d_g_* is the node degree of *g*, and the sum runs over all intermediate genes *g′* connecting *V* to *g*. For the application, we imposed a cut off *s* ≤ 0.001, which led to reasonable network sizes.

For any given function *F* from Table 1 we collect all viral proteins *V_i_* for which the intersection *G_i_* of the set of topscoring genes of *F* and the network neighborhood of *V_i_* is not empty, and together they form the “core” of the network for function *F*. Apart from the protein-protein binding edges, this core network can still be unconnected as a subgraph of the KG. In the next step, in order to show supporting evidence from the literature, we therefore include additional genes from the KG (with emphasis on high-scoring ones w.r.t. *F*) to create a connected subgraph. For the subset of gene nodes *g* in the KG that were assigned a *z*-score *z_gF_*, and that are connected by an edge, we define edge weights

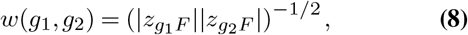

which are used to compute shortest paths from genes in the sets *G_i_* to the endpoint function *F* within the KG (similarly gene-function edges are assigned a weight *w* = |*z_gF_*|^−1/2^). Shortest paths constructed this way are enriched in high-scoring genes. In addition to shortest paths we also add slightly longer paths (based on a heuristic) to control network size. In a final step, in order to provide additional context, some network genes are also connected to nodes representing signaling pathways in which these genes play a role.

**Table 1.** Endpoint functions and pathways for which networks were computed. These networks represent the large spectrum of the host biology affected by the viral infection, including immunological signaling pathways, biological endpoints observed in severe or critically-ill COVID-19 patients, the viral life cycle and its counterpart host response, and host functions that are possibly hijacked by the virus itself for its replication/multiplication or transmission.

|  |  |  |
| --- | --- | --- |
| AMPK Signaling | Fragmentation of Golgi apparatus | Macroautophagy of cells |
| Antiviral response | Fusion of cells | Macropinocytosis |
| Apoptosis | Fusion of phagosomes | Mitosis |
| Asthma | Fusion of vesicles | Myocarditis |
| Autophagy | G Beta Gamma Signaling | PDGF Signaling |
| Budding of virus | Glycolysis of cells | PTEN Signaling |
| CDK5 Signaling | HIV infection | Phagocytosis |
| CXCR4 Signaling | HMGB1 Signaling | Pneumonia |
| Cell cycle progression | Hemoglobinopathy | Production of reactive oxygen species |
| Cellular homeostasis | Hemostasis | Pulmonary Hypertension |
| Chemotaxis | Hypertension | Release of virus |
| Clathrin mediated endocytosis | I-κB kinase/NF-κB cascade | Replication of RNA virus |
| Coagulation of blood | IGF-1 Signaling | Replication of coronavirus |
| Concentration of cholesterol | IL-1 Signaling | Respiratory failure |
| Degradation of Golgi apparatus | IL-2 Signaling | SAPK/JNK Signaling |
| Degranulation of cells | IL-6 Signaling | Stabilization of mRNA |
| Diabetes mellitus | IL-8 Signaling | Synthesis of lipid |
| Edema of pericardial cavity | Import of protein | Synthesis of phospholipid |
| Endocytosis | Infection by RNA virus | Synthesis of reactive oxygen species |
| Endoplasmic reticulum stress response of cells | Inflammation of respiratory system | Transport of virus |
| Engulfment of cells | Inhibition of ARE-Mediated mRNA Degradation Pathway | Viral Infection |
| Entrance of virus | Internalization of protein | Viral life cycle |
| Exocytosis | Interphase |  |
| Fibrosis | M phase |  |

## Discussion

Endpoint functions and pathways for which networks were computed are shown in Table 1. These networks represent the large spectrum of the host biology affected by the viral infection. Immunological signaling pathways (IL-1, IL-6, IL-8 to name a few) are included as they describe broadly the impact of the inflammatory setting in COVID-19 patients. We have also included networks that display biological endpoints observed in severe or critically-ill COVID-19 patients such as pneumonia, respiratory failure, myocarditis. A set of networks presents the complex viral life cycle and its counterpart host response (Replication of coronavirus, budding of virus, entrance of virus, antiviral response, etc.) and finally a set of networks that are possibly hijacked by the virus itself for its replication/multiplication or transmission (endocytosis, endoplasmic reticulum stress response, etc.). Below we discuss three of the networks as examples.

### Network: Promotion of coagulation of blood

Mostpeople infected with SARS-CoV-2 will experience mild to moderate respiratory illness and recover promptly. However, in some cases, severe disease occurs with major pathophysiological and sometimes lethal outcomes. Co-morbidities such as cardiovascular, renal, and respiratory preexisting conditions contribute to the severity seen in these patients. One drastic impact seen is the change in hemostasis after SARS-CoV-2 infection. Severe coagulopathy can arise and is associated with increased fatality rates in severely ill patients. The SARS-CoV-2 infection induces a pro-coagulative state and may result in vascular leakage and disseminated intravascular coagulation. The proinflammatory unbalance is thought to be one of the key factors of this uncontrolled clotting consequence seen in severe COVID-19 (3–7, 14). The network presented here (Figure 3) displays the interrelations between some of the key host molecules involved in the coagulation cascade, such as F3/tissue factor, F10, PLAT/plasminogen activator, and angiogenic factors or molecules related to angiogenesis balance such as F2RL1/proteinase-activated receptor 2 and EDN1/endothelin 1, PF4/Platelet factor 4 and several viral proteins (nsp9, nsp13, orf3a, orf9c, orf8). The severity of COVID-19 is generally a consequence of hyper-cytokinemia (“cytokine storm”) with its dramatic increase of chemokines and their cellular consequences (e.g. increase of neutrophils, thrombocytopenia, endothelialitis). Therefore, this network also displays the contribution of increased pro-inflammatory molecules or signaling pathways (IL1B, IL6, IL8, IL17a) and upregulated chemokine signaling (CXCR4 signaling, CCL5) observed in severe COVID-19 outcomes. This network highlights the importance of understanding the molecular interplay between the players, and as shown recently, anticoagulant treatment appears to decrease mortality in severe COVID-19 patients.

**Fig. 3.**
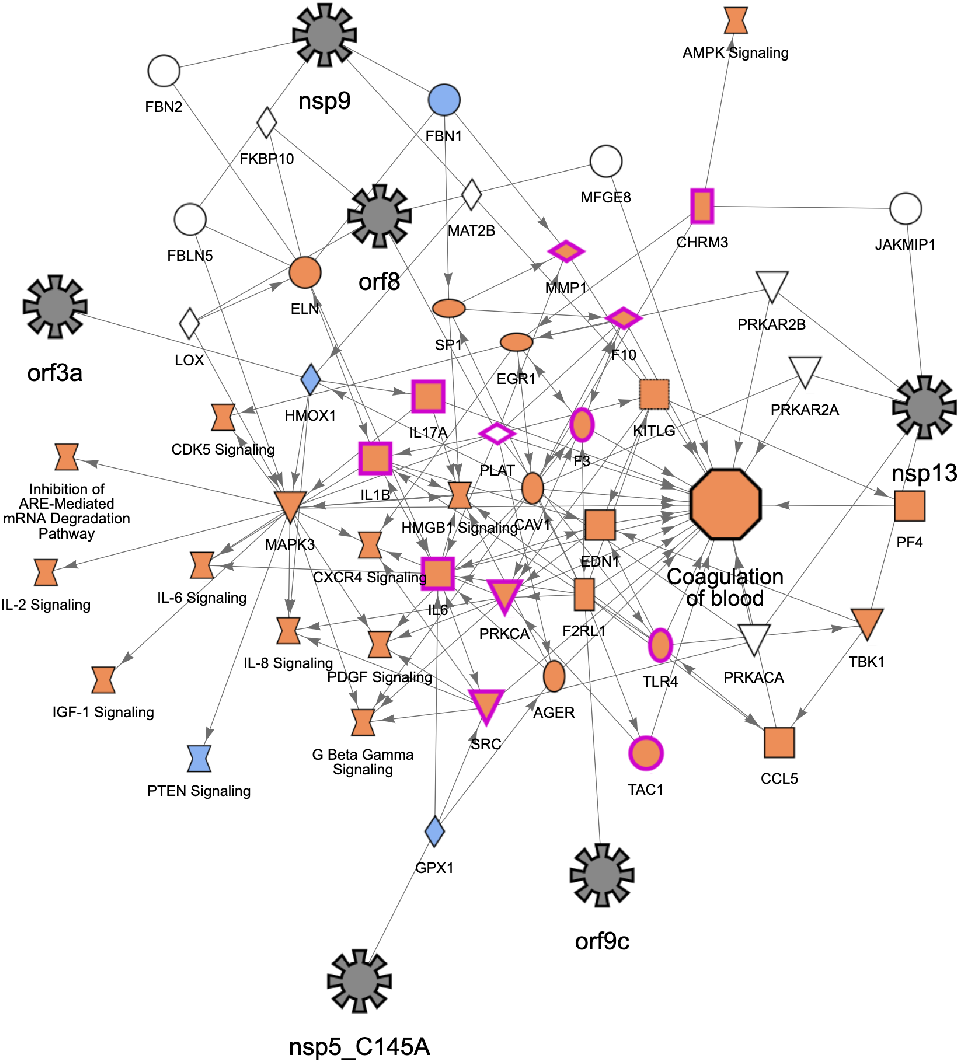
Network: Promotion of coagulation of blood. See Discussion section.

### Network: Promotion of pneumonia

Viral pneumonia with acute respiratory distress syndrome (ARDS) is one of the extreme consequences of COVID-19, a condition requiring mechanical ventilation as treatment. Patients with these severe conditions develop progressive respiratory failure following dramatic cascades of events. Dysfunctional immune responses in these patients will induce these events and are characterized by low IFN type I, III, a high pro-inflammatory setting, elevated chemokine secretion, high infiltration of myeloid and T cells in the lung, and finally severe pulmonary edema and pneumonia (15–17). This network (Figure 4) shows the possible interplay among type I Interferons, interleukins, the glucocorticoid receptor, sensors of viral infections and elements of the JAK/STAT pathway or the coagulation cascade and key coronaviral proteins that might promote pneumonia.

**Fig. 4.**
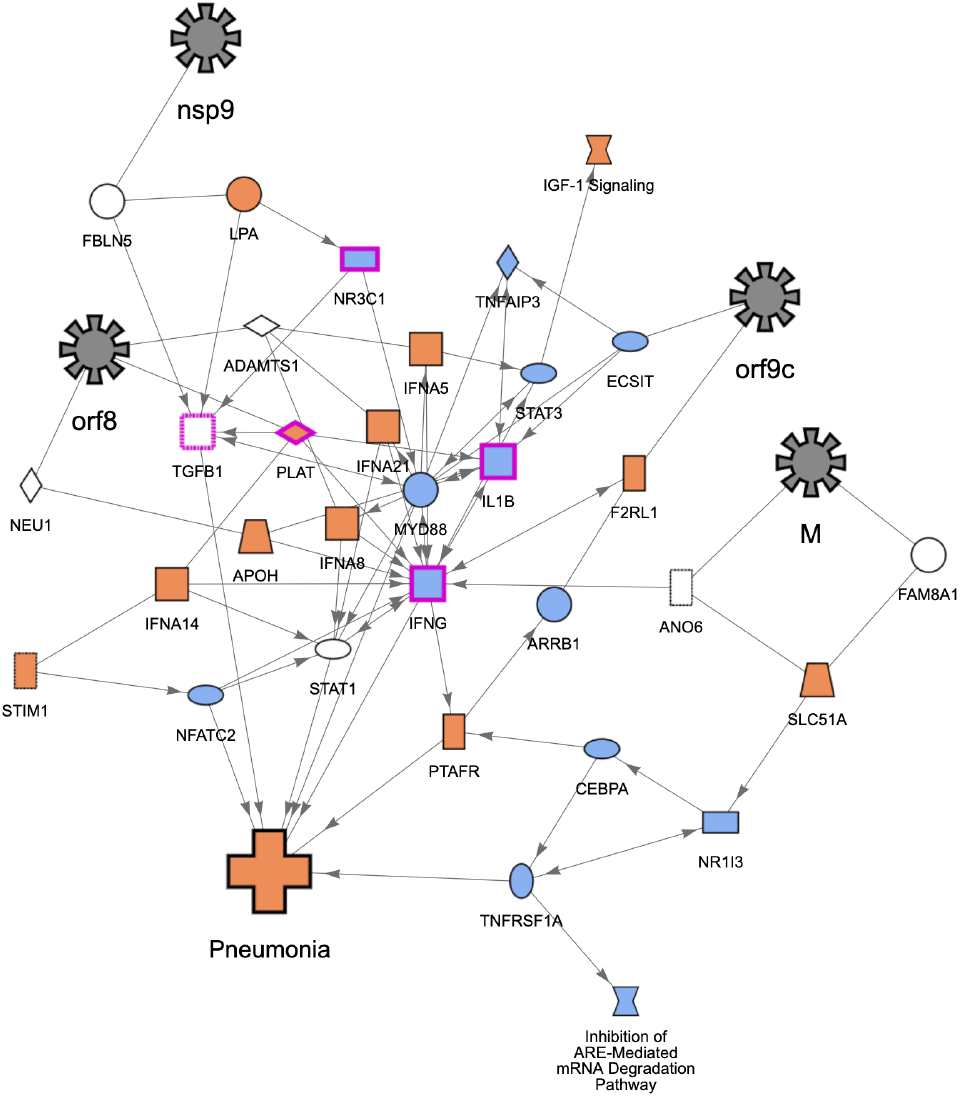
Network: Promotion of pneumonia. See Discussion section.

### Network: Promotion of IL6 signaling

IL6 is an important pleiotropic interleukin that signals through the JAK/STAT (JAK1 and STAT3 in particular) and the MAPK pathway. It is expressed by immune cells (dendritic cells, macrophages, B cells and also epithelial cells) and it is involved in many biological processes including cell survival, apoptosis, maturation of T cells, TH1/Th2/Th17 differentiation/balance, and inflammation. IL6 is described as a pro-inflammatory cytokine, secreted in response to IL1B and TNF stimulation. The severe or critical cases of COVID-19 correlate with high levels of IL6 (during the cytokine storm) and low lymphocyte counts (18–21). Furthermore, higher levels of IL6 correlate with the risk of developing ARDS. As such, clinical trials are now ongoing and are designed to target IL6 using monoclonal antibody therapy (Tocilizumab and Sarilumab) as IL6 seems to be one of the key promoters of fatal outcomes. IL6 signaling described in this network (Figure 5) includes all the major proinflammatory cytokines (IL1B, IL17, TNF and IL6,). Several key molecules of inflammation signaling are also present such as RELA, IKBKB, RIPK1 as well as MAPK signaling. It is thought that all these genes are either upregulated or predicted to be activated and would participate in the general increase of IL6 signaling and its unfortunate consequences in COVID-19.

**Fig. 5.**
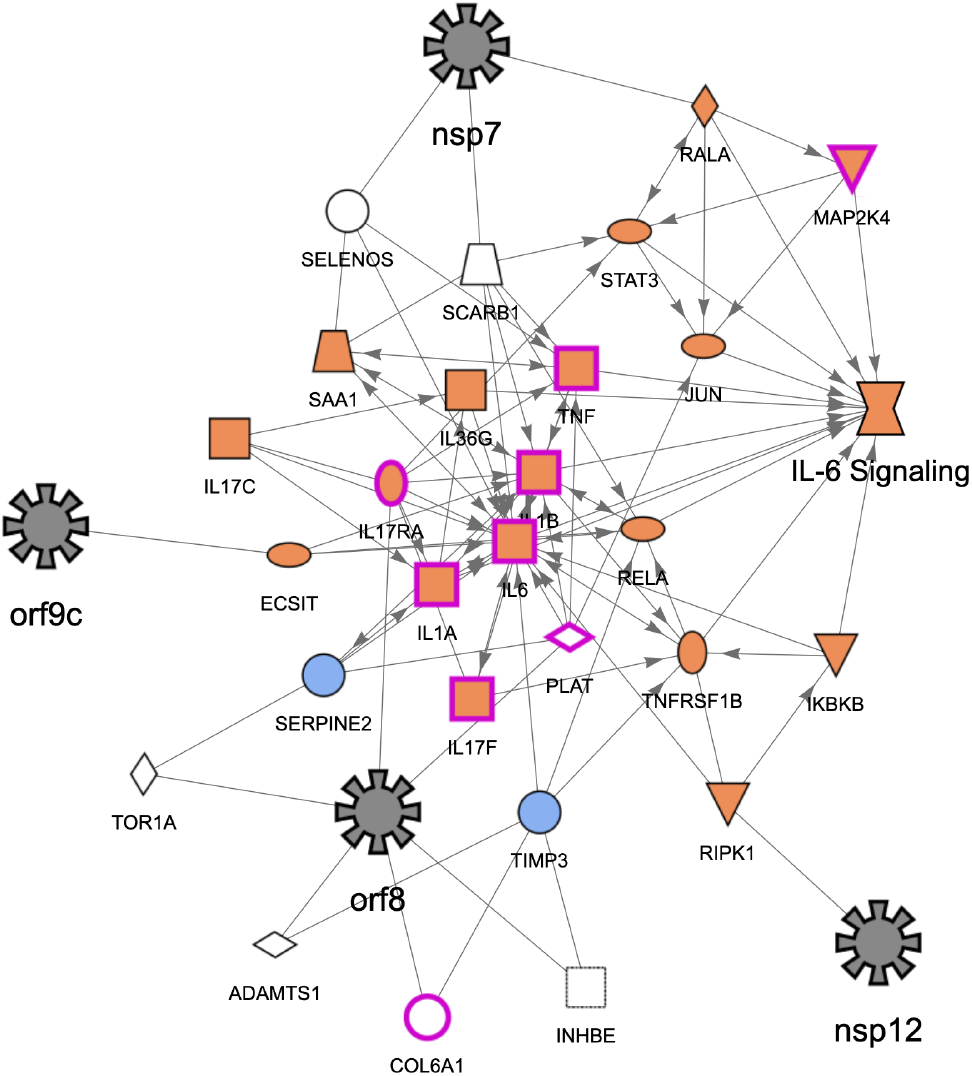
Network: Promotion of IL6 signaling. See Discussion section.

## Conclusion

We have presented the Coronavirus Network Explorer (CNE), a tool to explore how SARS-CoV2 may interfere with various cell functions through interactions with host genes. The CNE also suggests potential novel drug targets as well as existing drugs that could be repurposed against COVID-19. Our approach involves “mining” of a large-scale literature-derived knowledge graph by finding host genes that are predicted by a machine learning algorithm to be most relevant in a given functional context, and that are also closely associated with viral proteins through network interactions. These genes, connected viral proteins, and impacted biological pathways and processes are displayed together in a network in which edges generally represent experimentally observed relationships between nodes, illuminating potential underlying molecular mechanisms. We have discussed a small selection of these networks in order to demonstrate that our approach can identify biologically plausible hypotheses. These hypotheses are grounded in actual immunological, virological and pathological observations seen in SARS-CoV-2 infected patients as demonstrated by underlying literature findings. In future work, our plan is to further improve on these networks, making mechanisms more explicit by adding more relevant, possibly inferred causal connections, and taking into account specifc cell type or tissue contexts.

Interestingly, some of the drugs identified in our networks have been mentioned as potential therapeutic, or have been identified using other methods (22), validating our own approach. For instance, Baricitinib is a JAK1/2 inhibitor approved for Rheumatoid Arthritis and it is now tested in phases III clinical trial for treatment of moderate to severe COVID-19 patients (23, 24). Baricitinib’s mechanism proposal includes inhibition of viral entry via ACE2-mediated endocytosis. Our algorithm highlighted other potential mechanisms to counteract SARS-CoV-2 infection such as interfering with PDGF signaling in addition to the known inhibition of the gp130 cytokines (IL-6, IL-12, etc.). Anakinra, a known IL1-R antagonist has been also identified in two of our networks describing potential pathological consequences or co-morbidities factoring into COVID-19 (asthma, fibrosis). Anakinra is now tested in phases III clinical trials (25). Finally, we have also identified Siltuximab, an anti-IL6, present in 28 of 70 networks, this monoclonal antibody is currently being tested for potential COVID-19 treatment (26).

In conclusion, using a network algorithm combined with a machine learning model based on prior knowledge, we have explored possible molecular mechanisms of SARS-CoV-2 infection and proposed a set of networks that describe the pathogenesis of COVID-19. More validation will be required but evidence from on-going clinical trials confirm the well-foundedness of our proposal to discover new avenues for existing drugs.

## Acknowledgements

We would like to thank Bob Rebres and Andreea Pasare for guidance and discussion. We thank Shawn Bauer for critical reading of the manuscript, and especially thank Craig Upson for helping to build the CNE.

## Availability

The Coronavirus Network Explorer (CNE) has been made freely available to the scientific community as a web-based resource at https://digitalinsights.qiagen.com/coronavirus-network-explorer. Each interactive network has an individual URL that can be shared independently. In addition to the CNE, networks have also been made available within QIAGEN Ingenuity Pathway Analysis (IPA) (https://www.qiagen.com/qiagen-ipa) to allow exploration in the context of other datasets.

